# The Ndc80 complex is essential for the initial kinetochore-microtubule capture during early mitosis

**DOI:** 10.1101/2022.02.10.479964

**Authors:** Mohammed Abdullahel Amin, Destiny Ariel Wallace, Dileep Varma

## Abstract

Mitotic kinetochores are initially captured by dynamic microtubules via a ‘search-and-capture’ mechanism. The microtubule motor, dynein, is critical for kinetochore capture as it has been shown to transport microtubule-attached chromosomes towards the spindle pole during early mitosis. In metaphase, the kinetochore localized, microtubule-binding complex, Ndc80, plays a central role in stabilizing kinetochore-microtubule (kMT) attachments. It is not yet clear, however, if Ndc80, which is recruited to kinetochores very early during mitosis contributes to initial kMT capture. Here, by combining CRISPR/Cas9-mediated knockout and RNAi technology with assays specifically targeted to study kMT capture, we show that mitotic cells lacking Ndc80 exhibit severe defects in this function during prometaphase. Rescue experiments show that Ndc80 mutants deficient in microtubule-binding are unable to execute proper kMT capture. While cells inhibited of dynein alone are predominantly able to make initial kMT attachments, cells co-depleted of Ndc80 and dynein show severe defects in kMT capture. Further, we demonstrate a novel physical interaction between Ndc80 and dynein during prometaphase. Thus, our studies, for the first time, identify a distinct event in the formation of initial kMT attachments, which is directly mediated by Ndc80 followed by a coordinated function with dynein, both of which are required for efficient kMT capture and proper chromosome alignment.

## INTRODUCTION

Sister kinetochores are captured by the dynamic spindle microtubules during early mitosis to form kinetochore-microtubule (kMT) attachments, an essential pre-requisite for proper chromosome alignment and segregation. Kinetochores are large macromolecular machines assembled on the centromeric DNA that bridges sister chromatids to dynamic microtubules during mitosis and drive the accurate partitioning of the genetic material into the daughter cells. Over 100 proteins constitute the human kinetochores consisting of the inner kinetochore and the outer kinetochore regions (Cheeseman, 2014; Musacchio and Desai, 2017; Nagpal et al., 2015; Pesenti et al., 2016). While the inner kinetochore proteins connect with the centromeric DNA and form a platform for the assembly of the outer kinetochore, the outer kinetochore proteins are involved in microtubule-binding (Cheeseman and Desai, 2008; Santaguida and Musacchio, 2009). Even though a large number of outer kinetochore proteins have been shown to bind to microtubules, it is not clear which of these proteins actually contribute to the initial capture of human kinetochores during early stages of mitosis. Moreover, the mechanistic details of how kinetochore protein(s) enable the capture of kinetochores by microtubules during early mitosis is still not understood.

During early mitosis, outer surface of unattached kinetochores transiently expands outward to form a fibrous corona, and associate laterally with the sides of the microtubules (Barisic et al., 2014; Kapoor et al., 2006; Magidson et al., 2011; McEwen et al., 1998; Rattner and Bazett-Jones, 1989; Ris and Witt, 1981; Tanaka et al., 2005). The capture of chromosomes during early mitosis is primarily thought to be initiated as lateral contacts of kinetochores with the sides of spindle-pole-nucleated microtubules followed by the transport of captured kinetochores along the microtubule promoted by a kinesin-14 family member kar3 in yeast (Hayden et al., 1990; Tanaka et al., 2005). Previous work suggests that the minus-end directed microtubule motor, cytoplasmic dynein, is required for kinetochore capture, lateral attachment and poleward-motility in large cells such as newt pneumocytes (Rieder and Alexander, 1990). However, while succeeding work has supported dynein’s role in poleward motility via dynamic lateral contacts with spindle microtubules in smaller human cells in culture, it is not clear from these studies, the extent to which dynein function is required for the initial attachment of kinetochores to microtubules.

Among the other outer kinetochore proteins, the conserved Ndc80 complex consisting of Ndc80/Hec1, Nuf2, Spc24 and Spc25 plays a key role in forming robust interactions with microtubules during metaphase (DeLuca and Musacchio, 2012; Meraldi et al., 2006). The Ndc80 complex binds directly to microtubule polymers through a CH (Calponin Homology) domain and a positively charged, unstructured amino terminal tail (Cheeseman et al., 2006; Ciferri et al., 2005; Wei et al., 2007). High levels of phosphorylation of the tail domain by mitotic kinase, Aurora B, is thought to reduce the affinity of Ndc80 to microtubules in prometaphase, while lower phosphorylation levels in metaphase and anaphase enhances Ndc80 microtubule-binding to promote robust kMT attachments (Cheeseman and Desai, 2008; Deluca and Musacchio, 2012). Cells with disrupted Ndc80 complex function have severe defects in stabilizing kMT attachments in metaphase leading to extensive chromosome misalignment, mis-segregation (DeLuca et al., 2002; Desai et al., 2003), or even completely fail to segregate the chromosomes (Wigge et al., 1998; Wigge and Kilmartin, 2001). However, while the role of Ndc80 complex in the formation of stable kMT attachments has been studied intensively, a role in kMT attachment formation during early mitosis has not yet been attributed to this complex.

Based on the previous research in this area, we reasoned that the process of kMT capture during early mitosis could be further divided into two steps: (i) the initial kMT attachment formation and (ii) the ensuing poleward transport of kinetochores, which is known to be dependent on dynein. Both these steps need to be successfully completed to constitute a processive kinetochore capture event, which is in turn required for proper chromosome congression. The initial kMT attachments which are formed between dynamic microtubules and individual mono-oriented kinetochores are functionally distinct from the attachments that occur later in mitosis (during metaphase and anaphase), that is known to require the function of the Ndc80 complex. This is so because the initial kMT attachments are not load-bearing and are prone to easy detachments (to facilitate kinetochore motility on microtubules). On the other hand, the attachments in metaphase/anaphase are load-bearing (due to bi-attachment with microtubules of opposite spindle poles) and are stabilized with the help of protein factors that are accessory to Ndc80 (including the Ska1 complex, Cdt1, Astrin, etc.), to prevent kinetochore detachment during chromosome segregation (Amin et al., 2019; Cheeseman and Desai, 2008).

In this study, we specifically focus on the mechanism of the 1^st^ step in kMT capture, i.e., the formation of initial kMT attachments. By combining CRISPR/Cas9 gene editing and RNAi technology with targeted assays employing high-resolution confocal microscopy, we find that the function of the Ndc80 complex is essential for the formation of initial attachment kinetochore by microtubules during early mitosis. Further, we find that the expression of Ndc80 mutants deficient in microtubule attachment are unable to rescue the defects in kMT capture observed in Ndc80-depleted cells. We find that the inhibition of dynein however, only disrupted a substantially smaller fraction of initial kMT attachment events as compared to Ndc80 inhibition. Our studies point to a scenario where the dynein motor and the Ndc80 complex synergize during early mitosis to effect processive kinetochore capture.

## Results and Discussion

### Depletion of Ndc80 Complex Causes Defects in Initial Kinetochore-Microtubule Attachments

First, we systematically analyzed the localization of the microtubule-binding Hec1 subunit of the Ndc80 complex during early mitosis. Our immunofluorescence data show that Ndc80 is recruited to kinetochores during prophase, when no discernable dynein localization can yet be detected at kinetochores (Figure 1A, top panel) (Ciferri et al., 2005; DeLuca et al., 2005; Varma et al., 2013). Mitotic cells in prometaphase, however, showed strong localization of both dynein and Ndc80 to kinetochores (Figure 1A, bottom panel). We then disrupted the function of the Ndc80 complex either by depleting the Hec1 or Nuf2 subunit using RNAi technology or by knocking out the Nuf2 subunit using CRISPR/Cas9-mediated gene editing technology. Efficient loss of Ndc80 levels was validated by immunoblotting (Fig 1B) as well as immunostaining analyses (Fig S1A) for all the approaches we used for the functional perturbation of Ndc80.

**Figure 1.**
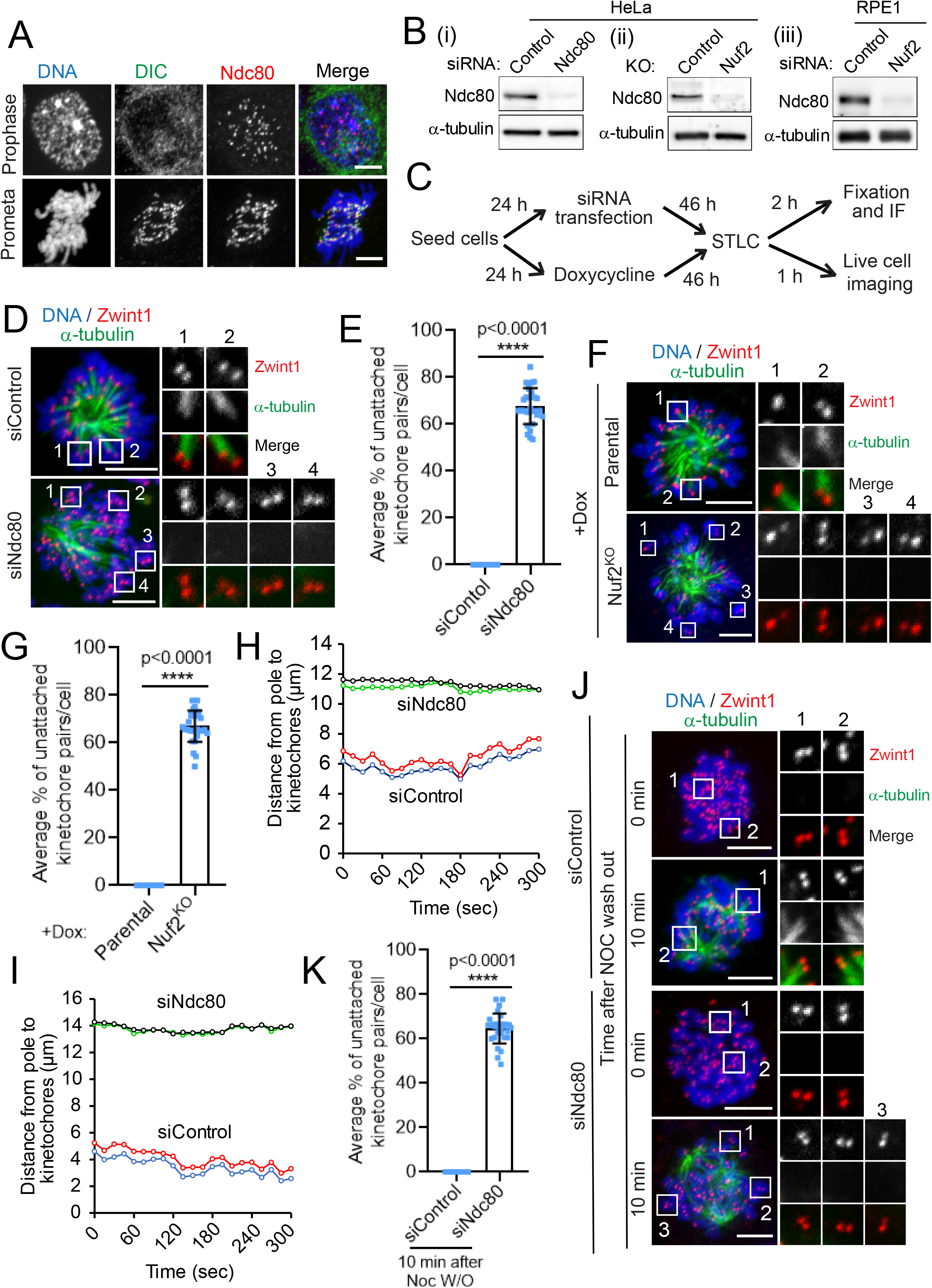
Initial kMT attachments are defective after the inhibition of the Ndc80 complex. (A) Immunofluorescence staining of dynein intermediate chain (DIC, green), and Ndc80 (red), in early mitotic prophase (top panels) or prometaphase (bottom panel), with the chromosomes counterstained using DAPI. Bars, 5 μm. (B) Western blot analysis of HeLa cells treated with siRNAs for control and Ndc80 (Bi), of Hela cells of parental and Nuf2 CRIPR/Cas9 KO (Bii), and of RPE1 cells treated with siRNAs for control and Nuf2 (Biii). α-tubulin was used as a loading control. (C) Cells were subjected to the above-mentioned perturbations, then treated with the indicated drugs/chemicals and fixed according to the scheme. (D and F) Immunofluorescence staining of STLC treated mitotic HeLa cells depleted of Ndc80 (D) or knocked out of Nuf2 (F) in comparison to respective control^RNAi^ or parental control cells, and stained for α-tubulin (green), a kinetochore marker Zwint1 (red) with the chromosomes counterstained using DAPI. Bars, 5 μm. Inset shows the kMT attachment status of individual kinetochore pairs in the conditions indicated. (E and G) Quantification of the status of kMT attachments in cells from D and F. Error bars represent S.D. from three independent experiments. For each experiment, on average ~ 20 kinetochore pairs from 10 different monopolar cells were examined. ****P < 0.0001 (Student’s t test). (H and I) Trajectories of two representative sister kinetochore pairs in HeLa cells transfected with siRNAs for control and Ndc80. 50 kinetochore pairs were analyzed for each condition. Also see Supplemental Videos 1 and 2. (J) HeLa cells treated with control^RNAi^ or Ndc80^RNAi^ were fixed 0 or 10 min after nocodazole washout. The cells were then immunostained for α-tubulin (green), a kinetochore marker Zwint1 (red), with chromosomes counterstained using DAPI. Bars, 5 μm. Inset shows the kMT attachment status of individual kinetochore pairs at these two time points under either condition. Also see Figure S2A. (K) Quantification of the status of kMT attachments in cells from J. Error bars represent S.D. from three independent experiments. For each experiment, 10 mitotic cells were examined. ****P < 0.0001 (Student’s t test).

To accurately test for the role of Ndc80 in initial kMT attachment step, we choose to first resort to a method that simulates the kMT attachments formed in early mitosis during the ‘search and capture’ process that are mono-oriented and not load-bearing in nature. We treated cells with a small molecule S-Trityl-L-Cysteine (STLC), which inhibits the kinesin, Eg5, and mitotic spindle bipolarity to produce monopolar spindles (Fig 1C) (Debonis et al., 2008; Mayer et al., 1999). We observed that STLC-treated monopolar cells depleted of the Hec1 subunit of the Ndc80 complex (Ndc80^RNAi^) had significantly higher number of unattached kinetochore pairs compared to that of cells treated with control siRNA (control^RNAi^) and STLC. Careful analysis revealed that as compared to control cells where almost all kinetochore pairs had at least one kinetochore attached to the microtubules, ~ 70% kinetochore pairs in Ndc80^RNAi^ cells were completely unattached, among all discernable kinetochore microtubule interaction events analyzed (Fig 1D, E). We also observed that STLC-treated HeLa cells knocked out of Nuf2 (Nuf2^KO^) had a similar, significantly higher fraction (> 65%) of kinetochore pairs which do not attach to microtubules as compared to that of parental cells (Fig 1F, G). Similar defects in initial kMT attachments (~ 65% unattached) were observed in diploid RPE1 cells treated with STLC and depleted of Nuf2 (Nuf2^RNAi^) as compared to control^RNAi^ cells (Fig S1B, C). To further examine the initial kMT attachment events, we analyzed the dynamic oscillation of sister kinetochore pairs in STLC-treated cells by live cell imaging. Our live imaging data showed that kinetochores in control^RNAi^ cells have robust oscillatory motion as expected as they were attached properly to the ends of dynamic kMTs (Fig 1H, I, siControl traces, video S1). In contrast, kinetochores in Ndc80^RNAi^ cells showed remarkably lower incidence of oscillatory motion, undergoing very short-range, erratic or random, back-and-forth motion instead (Fig 1H, I, siNdc80 traces, video S2).

### Other Kinetochore Proteins Required for Proper Chromosome Alignment are not Critical for Initial Kinetochore-microtubule Attachment Events

As chromosomes are captured at the interface of outer kinetochores with microtubules, we next investigated the role of other outer microtubule-binding and accessory kinetochore proteins known to have a role in proper chromosome alignment, including the dynein-dynactin complex, CLIP-170, Spindly and CENP-E (Amin et al., 2014; Galjart, 2005; Gassmann et al., 2008; Huang et al., 2012; Kardon and Vale, 2009; Yu et al., 2019). Dynein, CLIP-170 and CENP-E have been proposed to generate lateral interactions between kinetochores and microtubules to aid in bi-directional chromosome movements during their congression to the metaphase plate (Sardar and Gilbert, 2012; Tanenbaum et al., 2006; Vorozhko et al., 2008). Spindly has been reported to be involved in rapid integration of peripheral chromosomes into the mitotic spindle but not as such in chromosome attachment to microtubules (Barisic et al., 2010). Moreover, Spindly is involved in the prevention of load-bearing attachment formation during prometaphase in *C. elegans* (Gassmann et al., 2008). We first confirmed the localization to these proteins to prometaphase kinetochores during early mitosis (Fig S2A). Next, we disrupted their function by treating with specific siRNAs and employing published protocols (Amin et al., 2015; Amin et al., 2018; Kiyomitsu et al., 2007). We confirmed the efficiency of depletion of the target proteins by immunoblotting (Fig S2B). We then tested if these outer kinetochore proteins have a role in initial kMT attachments by carefully visualizing these attachment events using the same STLC-treatment assay as we did for the Ndc80 inhibition experiments. Our data show that initial kMT attachments in STLC-treated cells depleted of dynein, p150glued (a dynactin subunit), CLIP-170, Spindly, CENP-E or Knl1 was surprisingly normal as observed in control^RNAi^ cells (Fig S2C, S2D). However, in cells depleted of all of the above-mentioned proteins (especially dynein and its accessory proteins), the chromosomes remained tethered to the plus-ends of microtubules constituting the monopolar spindles, consistent with previous work (Varma et al., 2008). These data suggest that most of the outer kinetochore proteins we tested, apart from the Ndc80 complex do not play an essential role in forming initial kMT attachments during early mitosis but rather could be involved in other mitotic functions, including chromosome congression or the spindle assembly checkpoint. As described earlier, the inhibition of the Ndc80 complex on the other hand, severely compromised the initial step in kMT capture, i.e., the formation of initial kMT attachments, yielding the more severe phenotype in defective kMT capture.

Our data supports the previous observations that dynein motor activity is critical for the 2^nd^ step in the kMT capture process, i.e. the poleward motility of the kinetochores, which ensues immediately after the formation of initial kMT attachments during early mitosis (Rieder and Alexander, 1990; Varma et al., 2008; Vorozhko et al., 2008). Dynein-mediated lateral contacts are currently also understood to be the primary mode of initial kMT attachment formation. However, our observations along with the ability of the Ndc80 complex to facilitate load-bearing, end-on kMT attachments in metaphase that is well-established, suggest that a considerable fraction of initial kMT attachment events in cultured human cells during early mitosis could also possibly be an outcome of dynamic end-on contact of microtubule plus-ends with individual kinetochores during the ‘search and capture’ process, rather than predominantly via lateral contacts as is commonly conceived. We propose that these dynamic end-on kMT attachments during early mitosis are mediated by the activity of the Ndc80 complex, but with the key distinction that these attachments are not stabilized (and the kinetochore pairs are not bioriented) as in metaphase cells. Our findings also support the observations from previous studies that only minor defects in chromosome congression is observed in dynein or CENP-E depleted cells (Kapoor et al., 2006; Putkey et al., 2002; Varma et al., 2008; Yang et al., 2007), where only dynamic lateral initial kMT contacts are prevented. We predict that these lateral attachments, mediated by the above-mentioned kinetochore motors, only account for ~ 30% of all initial attachment events. Further, our results suggest that the loss of function of these kinetochore motor proteins could be compensated for by the activity of the Ndc80 complex over time to produce nearly normal chromosome alignment. Future live imaging studies of early mitotic cells with higher optical and temporal resolution are required to determine the fraction of initial kMT attachments facilitated by the Ndc80 complex that are either dynamic lateral or dynamic end-on in nature.

### *De novo* Microtubules do not Attach to Kinetochores in Ndc80-depleted Cells

*De novo* microtubules can be generated by treating cells with nocodazole to completely disassemble the cellular microtubules followed by washing out the drug and releasing into fresh culture medium (Cimini et al., 2001; Piperno et al., 1987). We further investigated the role of Ndc80 in the initial attachment of kinetochores to *de novo* microtubules using the nocodazole washout assay (Fig S3A). Nocodazole completely disassembles all the cellular microtubules; consequently, we found that HeLa cells treated either with Ndc80^RNAi^ (Fig 1J, K) or subjected to Nuf2^KO^ (Fig S3B, C) had no cellular microtubules at zero mins after nocodazole washout as compared to their respective controls. After 10 minutes of nocodazole washout, most of the kinetochores in control^RNAi^ HeLa cells or control parental HeLa cells were found to be attached to *de novo* microtubules. We insinuate that these microtubules could either be emanating from the centrosome (to make contact with the kinetochores later) or could directly be originating from the kinetochores themselves (Maiato et al., 2004; Tulu et al., 2006). In contrast, a significant number (~ 65%) of kinetochores were found to be not attached to *de novo* microtubules in Ndc80^RNAi^ or Nuf2^KO^ cells (Fig 1J, K; Fig S3B, C). We also observed similar defects in initial kMT attachments in Nuf2^RNAi^ RPE1 cells as compared to control^RNAi^ cells. Nuf2^RNAi^ RPE1 cells had a significant number (~ 65%) of kinetochores, which were unattached to the *de novo* microtubules 10 mins after nocodazole washout compared to that in control^RNAi^ cells (Fig S3D, E). These data further indicate that kinetochores are not able to attach to newly formed microtubules in the absence of the Ndc80 complex during early mitosis. Interestingly, these experiments also suggest that the Ndc80 complex could contribute to the formation of initial kMT attachments by two mechanisms. First, the function of the complex could be required for directly forming lateral or end-on dynamic contacts of kinetochores with spindle-pole focused microtubules. Secondly, the microtubule-binding activity of Ndc80 could be critical for *de novo* kMT formation from kinetochores.

### Depletion of Ndc80 Complex Causes Defects in the Formation of Initial Kinetochore-Microtubule Attachments in Physiological Conditions

Having already employed two distinct assay conditions to monitor initial kMT attachment formation in Ndc80-inhibited cells, we next tested the status of initial kMT capture under normal physiological conditions by using both fixed and live cell imaging (Fig 2A). Our fixed cell analysis showed that Ndc80^RNAi^ HeLa cells had significantly higher number (> 65%) of kinetochore pairs which are unattached to microtubules during early mitosis compared to that of control^RNAi^ cells (Fig 2B, C). We measured the distance of separation between kinetochore and the nearest microtubule loci (attached or unattached) to better define the status of the initial kMT attachments. We observed that the average kinetochore-microtubule distance was ~ 85 nm when kinetochores are attached to microtubules in control^RNAi^ cells. On the other hand, in Ndc80^RNAi^ cells, while there were no microtubules for distances > 1 μm within the vicinity for many kinetochores, the average kinetochore-microtubule distance was found to be ~ 750 nm in cases where a microtubule was observed to be in the vicinity of a kinetochore (Fig. 2E). We found that Nuf2^KO^ HeLa cells also had > 65% of their kinetochore pairs not captured by microtubules in early mitosis as compared to that of parental cells (Fig 2B, D). Besides HeLa cells, we observed similar defects (~65%) in initial kMT attachments in Nuf2^RNAi^ RPE1 cells (Fig 2F, G).

**Figure 2.**
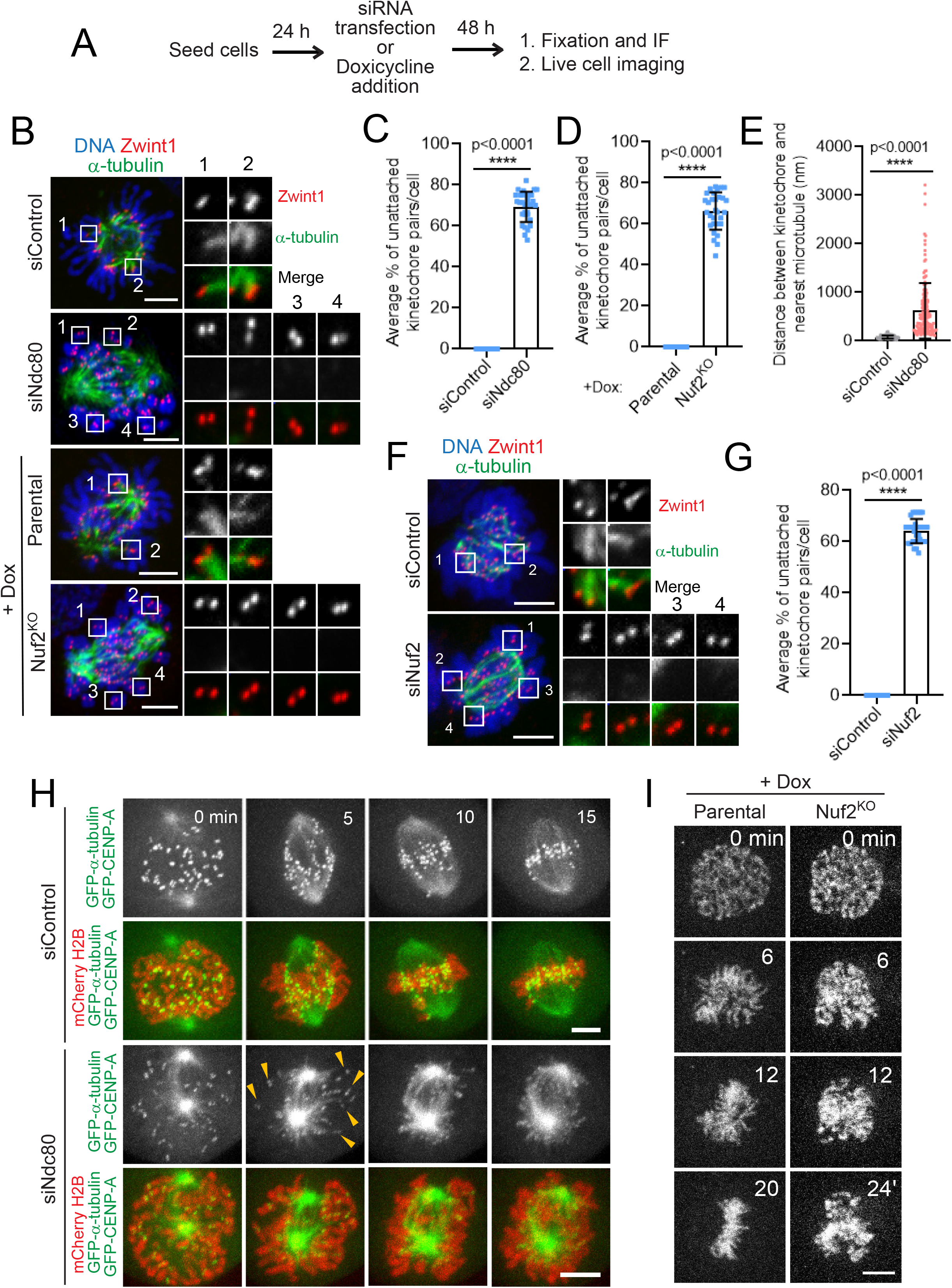
Role of Ndc80 complex in kMT capture during early mitosis examined using fixed and live cell imaging under physiological conditions. (A) Cells were subjected to the indicated perturbations, treated with the indicated drugs/chemicals and fixed according to the scheme. (B) Immunofluorescence staining of early prometaphase HeLa cells depleted of Ndc80 (top two panels) or knocked out of Nuf2 (bottom two panels) in comparison to control^RNAi^ or parental cells respectively and stained for α-tubulin (green) and a kinetochore marker, Zwint1 (red) with the chromosomes counterstained using DAPI. Bars, 5 μm. Inset shows the kMT attachment status of individual kinetochore pairs in the indicated conditions. (C and D) Quantification of the status of kMT attachments in cells from B. Error bars represent S.D. from three independent experiments. For each experiment, on average ~20 kinetochore pairs from 10 early prometaphase cells were examined. ****P < 0.0001 (Student’s t test). (E) The distance measured between kinetochore and nearest microtubule in cells from B. Error bars represent S.D. from three independent experiments. For each experiment, on average 20 kinetochore pairs from 5 early prometaphase cells were examined. ****P < 0.0001 (Student’s t test). (F) Immunofluorescence staining of early prometaphase RPE1 cells depleted of Nuf2 in comparison to control^RNAi^ cells (top panel) and stained for α-tubulin (green), a kinetochore marker Zwint1 (red) with the chromosomes counterstained using DAPI. Bars, 5 μm. Inset shows the kMT attachment status of individual kinetochore pairs. (G) Quantification of the status of kMT attachments in cells from F. Error bars represent S.D. from three independent experiments. For each experiment, on average ~20 kinetochore pairs from 10 early prometaphase cells were examined. ****P < 0.0001 (Student’s t test). (H) Selected frames from live imaging of double-thymidine synchronized HeLa cells stably expressing mCherry-Histone H2B to visualize the chromosomes, in addition to GFP-α-tubulin and GFP-CENPA to visualize kMT attachment and treated with control^RNAi^ (top two panels) or Ndc80^RNAi^ cells (bottom two panels). Images were captured every 1 min interval for 15-20 min until chromosome is aligned in control^RNAi^. Bars, 5 μm. Yellow arrow heads indicate the kinetochores unattached to microtubule in Ndc80^RNAi^ cells. The images from the movies were cropped and image intensity adjusted as required for better visualization of kinetochores and microtubules. (I) Selected frames from live imaging of parental (left panel) and Nuf2^KO^ (right panel) HeLa cells treated with Hoechst for 30 min prior to imaging to visualize the chromosomes. Images were captured every 2 min interval for around 30 min until all the chromosomes were aligned in parental cells. Bars, 5 μm.

We further confirmed the defects in initial kMT attachments in Ndc80-inhibited cells under normal physiological conditions by observing kMT behavior during early mitosis in live cells. Previous studies using both fixed and live-imaging experiments (Amin and Varma, 2017; Miller et al., 2008; Sundin et al., 2011) suggest that mitotic cells are unable to align a large fraction of their chromosomes for an extended period of time after Ndc80 inhibition. Apart for visualizing kMT behavior, the labelling of spindle microtubule in our live experiments also ensured that bipolar spindles are formed normally in Ndc80-inhibited cells. To ensure that the observed phenotype was not accumulated over several cell cycles, we chose to use double-thymidine synchronized HeLa cells, where Ndc80 knockdown was performed only during the two successive periods of thymidine treatment, for our live imaging (Amin and Varma, 2017; Varma et al., 2012). Our studies show that in control^RNAi^ cells, all the kinetochores are attached to microtubules within 5-10 mins after nuclear envelope breakdown (NEB) and are completely aligned at the metaphase plate within 15-20 mins (Fig 2H, top two panels, Video S3). In contrast, a substantial number of sister kinetochore pairs (~65%) of Ndc80^RNAi^ cells remained unattached to microtubules and were not able to align properly at the metaphase plate in the same duration of time (Fig 2H, bottom two panels, Video S4). Live cell imaging of Nuf2^KO^ HeLa cells with their DNA labelled using Hoechst further shows that chromosome alignment is severely impeded in this scenario as compared to parental controls supporting the live-imaging results obtained with Ndc80^RNAi^ synchronized HeLa cells (Fig 2I, Video S5 and 6).

### Ndc80’s Role in Initial Kinetochore-Microtubule Attachment is Directly Dependent on its Ability to Bind Microtubules

To obtain additional insight into the role played by Ndc80 in initial kMT attachments, we performed rescue experiments using different Ndc80 mutants that have been shown to be defective in forming load-bearing kMT attachments in metaphase. The microtubule-binding activity of the Ndc80 complex is known to be located at two distinct elements within the amino terminus of its Hec1 subunit: a positively charged, unstructured amino terminal tail with Aurora B phosphorylation sites dispersed within this region, and the adjacent CH domain, a globular microtubule-binding domain found in proteins (Fig. 3B) (Cheeseman et al., 2006; Ciferri et al., 2008; DeLuca et al., 2006; Maskell et al., 2010; Miller et al., 2008; Petrovic et al., 2010). High-resolution cryo-EM and *in vitro* analyses has further shown that both the N-terminal tail and CH domains are important for efficiently binding to microtubules (Alushin et al., 2012; Alushin et al., 2010; Lampert et al., 2013; Sundin et al., 2011; Tooley et al., 2011; Wilson-Kubalek et al., 2008) (Ciferri et al., 2008; Lampert et al., 2013; Sundin et al., 2011; Tooley et al., 2011; Wei et al., 2007). In cells, the N-terminal tail and CH domain of Hec1 have been reported to be critical to form stable kMT attachments (Guimaraes et al., 2008; Miller et al., 2008; Sundin et al., 2011). Rescue experiments showed that the stability of kMT attachments is affected by the phosphorylation state of the N-terminal tail domain of Hec1. Cells expressing a nonphosphorylatable Hec1 tail domain exhibited hyperstable kMT attachments while cells expressing phosphomimetic Hec1 tail mutant had unstable kMT attachments (DeLuca et al., 2006; Guimaraes et al., 2008; Welburn et al., 2010). We thus tested whether the N-terminal phosphomutants of Hec1 contribute to its role in initial kMT attachment using the indicated experimental scheme (Fig. 3A). Our rescue experiments showed that Ndc80^RNAi^ cells transfected with Hec1^WT^ or nonphorylatable Hec1^9A^ mutant constructs and treated with STLC had no defects in the formation of early kMT attachments as expected (Fig 3C, D). In contrast, Ndc80^RNAi^ cells transfected with phosphomimetic Hec1^9D^ mutant and treated with STLC had a significant fraction (~65%) of kinetochores which were not attached to microtubules (Fig 3C, D). These data suggest that initial kMT attachments are directly influenced by the Aurora B kinase-mediated phosphorylation of the N-terminal tail of Ndc80.

**Figure 3.**
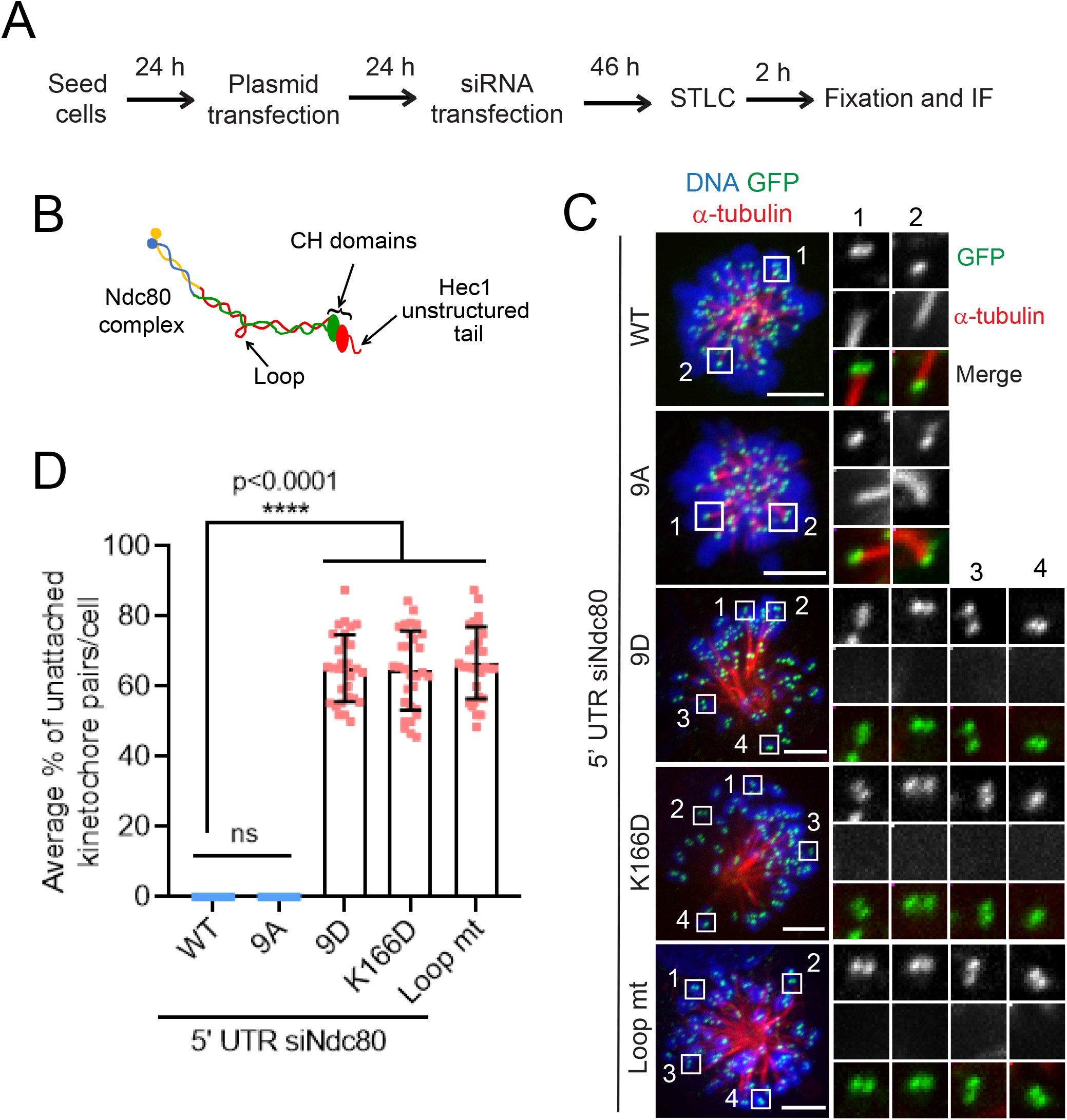
Characterization of different microtubule-binding domains within the Ndc80 complex required for the initial kMT attachments. (A) Cells were subjected to the indicated perturbations, treated with the indicated drugs/chemicals, and fixed according to the scheme. (B) A cartoon showing the Ndc80 complex with different domains of the Hec1 subunit that have been implicated in kMT attachments. (C) Immunofluorescence staining of STLC-treated mitotic HeLa cells depleted of endogenous Ndc80 and transfected with different Hec1-GFP constructs as indicated and stained for GFP (green), α-tubulin (red) with the chromosomes counterstained using DAPI. Bars, 5 μm. Inset shows the kinetochore microtubule (kMT) attachment status of individual kinetochore pairs in the conditions indicated. (D) Quantification of the status of kMT attachments in cells from C. Error bars represent S.D. from three independent experiments. For each experiment, on average ~20 kinetochore pairs from 10 monopolar cells were examined. ****P < 0.0001 (Student’s t test).

As described earlier, mutations in its CH domain have been reported to severely impair Ndc80’s microtubule-binding *in vitro* and the formation of stable kMT attachment *in vivo* due to a change in protein conformation that compromises this activity (Ciferri et al., 2008; Lampert et al., 2013; Sundin et al., 2011; Tooley et al., 2011). Thus, we also tested whether role of Ndc80 in initial kMT capture is influenced by the CH domain mutant (Hec1^K166D^), defective in microtubule-binding. Our rescue experiments showed that Ndc80^RNAi^ cells transfected with the Hec1^K166D^ mutant construct and treated with STLC also had a significant fraction (~65%) of kinetochores which do not attach to microtubules as compared to cells rescued with Hec1^WT^ (Fig 3C, D). These data suggest that CH domain-mediated physiological conformation of Hec1 is also essential for in initial kMT capture.

In addition to the established microtubule-binding sites within the Ndc80 complex, the internally located loop domain has also been reported to be important for the recruitment of Ndc80 accessory proteins such as the Ska complex and Cdt1 to human kinetochores, where they are required to form robust kMT attachments during metaphase (Amin et al., 2019; Monda and Cheeseman, 2018). We hence used the STLC treatment assay to test if the Hec1 loop domain was required for kMT capture. Surprisingly, we find that rescue experiments with a Hec1 loop domain mutant (Hec1^LoopMut^) also exhibited similar defects in kinetochore capture as observed with the other mutants defective in direct microtubule-binding (Fig. 3C, D). Further studies are required to define the mechanism by which the loop domain contributes to initial kMT attachments during early mitosis.

### Depletion of the Ndc80 Complex Leads to Kinetochore-null Phenotype When Aurora B Kinase is Inhibited Simultaneously

kMT attachments are regulated by Aurora B kinase-mediated variable phosphorylation of the Ndc80 complex during the different stages of mitosis (Cheeseman et al., 2006; DeLuca et al., 2006; Murata-Hori et al., 2002). While Aurora B destabilizes kMT contacts during early mitosis to aid in attachment error correction, inhibition of the kinase activity using small-molecule inhibitors leads to premature stabilization of kMT attachments of non-bioriented prometaphase kinetochores (Hauf et al., 2003; Lampson et al., 2004; Tanaka et al., 2002). We thus reasoned that hampering Aurora B activity using the single molecule inhibitor, ZM447439, (using the indicated protocol, Fig 4A) should rescue the defects in initial kMT attachments observed in Ndc80-inhibited cells during early mitosis. For our experiments, we coupled the inhibition of Aurora B with MG132 treatment to prevent premature anaphase onset. We found that control^RNAi^ or parental mitotic HeLa cells attained partial chromosome alignment, presumably because the formation of initial kMT attachments occurred normally as expected in these cells (Fig 4B, 4C, top panels, Fig 4D). Surprisingly, after ZM447439 treatment, kinetochores of around 75% mitotic Ndc80^RNAi^ or Nuf2^KO^ cells were found completely outside of the mitotic spindle, uncoupled from the microtubules, a phenomenon previously referred to as “kinetochore-null” phenotype (Gassmann et al., 2008; Oegema et al., 2001) (Fig 4B, 4C bottom panels, Fig 4D). We then asked whether the kinetochore-null phenotype is produced under similar conditions but even in the presence of paclitexal (Taxol), a small molecule that stabilizes microtubules in general, thus also serving to rescue the combined effects of Aurora B and Ndc80 inhibition, at least partially. We found that a significant number of mitotic Ndc80^RNAi^ or Nuf2^KO^ (Fig S4A, B, bottom panels; Fig S4C) cells continued to exhibit the kinetochore-null phenotype even in this scenario (ZM447439 + Taxol) as compared to control^RNAi^ cells or parental cells (Fig S4A, B, top panels; Fig S4C). Further, as observed in HeLa cells, 75% of mitotic Nuf2^RNAi^ RPE1 cells also showed kinetochore-null phenotype which was significantly higher as compared to that of control^RNAi^ cells after the inhibition of Aurora B (Fig 4E, F). We then analyzed this interesting kinetochore-null phenotype using live cell imaging. No control^RNAi^ cells showed this phenotype in the presence of ZM447439 during the period (~ 25 mins) of our live imaging, suggesting that kinetochore capture, and alignment is normal in this case (Fig 4G, H, top two panels, video S7). In contrast, a significantly higher portion (~ 70%) of mitotic Ndc80^RNAi^ cells showed the kinetochore-null phenotype (Fig 4G, H, bottom two panels, video S8) where kinetochores (marked by GFP-fused CENP-A) were observed to remain completely separated away from the vicinity of the mitotic spindle during the entire period of our live imaging procedure, without initiating any capture events.

**Figure 4.**
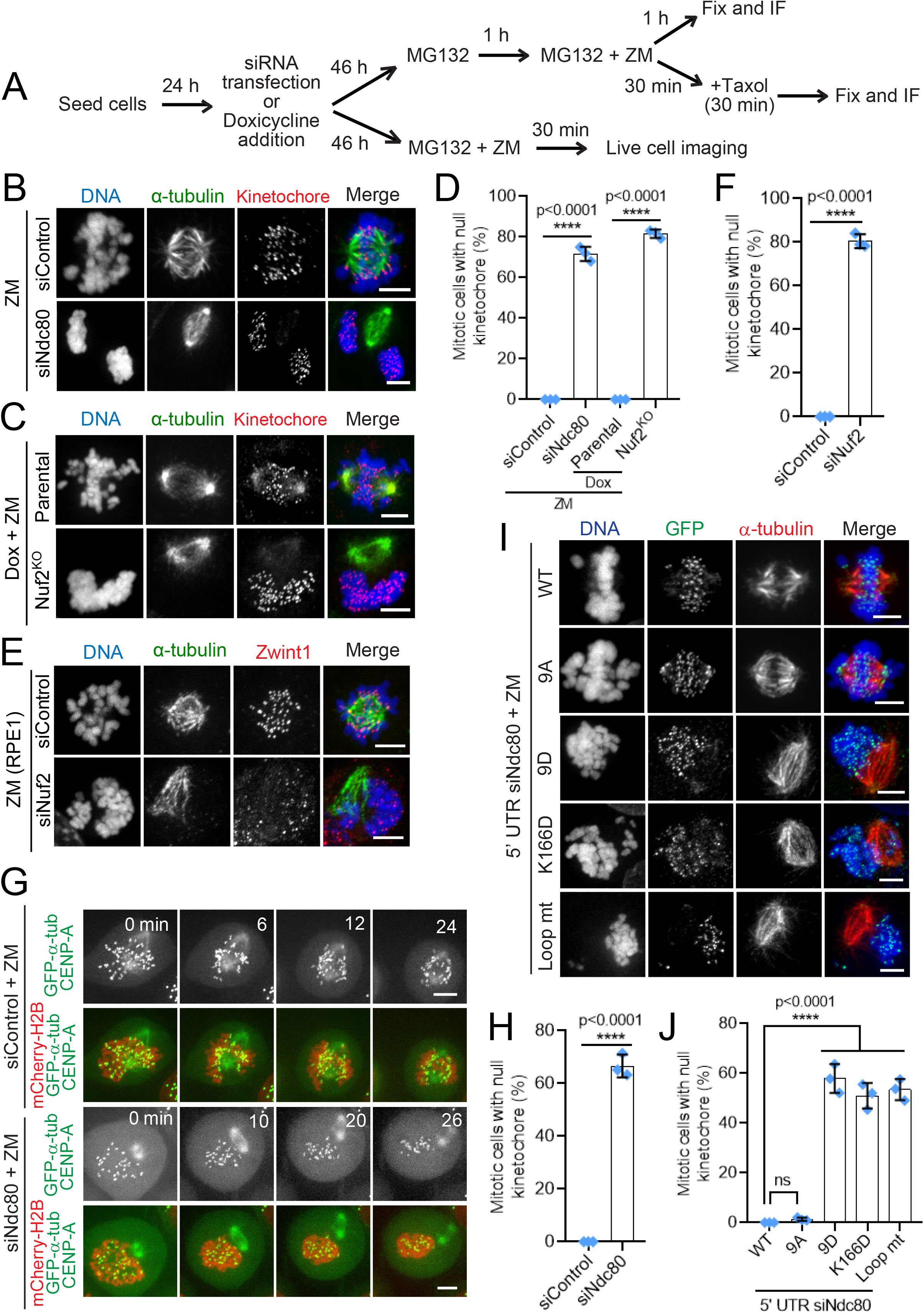
Characterization of the kinetochore-null phenotype induced by Ndc80-inhibition. (A) Cells were subjected to the indicated perturbations, treated with the indicated drugs/chemicals, and fixed according to the scheme. (B and C) Mitotic HeLa cells depleted of Ndc80 (B, panel) or knocked out of Nuf2 (C bottom panel) as compared to control^RNAi^ (B, top panel) or parental control cells (B, bottom panel), and followed by the indicated drug treatments, were immunostained for α-tubulin (green), a kinetochore marker Zwint1 or CENPA (red) with the chromosomes counterstained using DAPI. Bars, 5 μm. (D) Quantification of the frequency of mitotic cells with null kinetochores (see main text for more details) in samples from B and D. Error bars represent S.D. from three independent experiments. For each experiment, 200 mitotic cells were examined. ****P < 0.0001 (Student’s t test). (E) Immunofluorescence staining of mitotic RPE1 cells depleted of Nuf2 in comparison to control cells (top panel) and stained for α-tubulin (green), a kinetochore marker Zwint1 (red) with the chromosomes counterstained using DAPI. Bars, 5 μm. (F) Quantification of the status of kMT attachments in cells from E. Error bars represent S.D. from three independent experiments. For each experiment, 200 mitotic cells were examined. ****P < 0.0001 (Student’s t test). (G) Selected frames from live imaging of HeLa cells stably expressing mCherry-Histone H2B to visualize the chromosomes in addition to GFP-α-tubulin and GFP-CENPA to visualize kMT attachment in control^RNAi^ (top two panels) and Ndc80^RNAi^ cells (bottom two panels). Images were captured at every 1 min interval starting from the point of nuclear envelope breakdown for around 30 min until a proper mitotic spindle was formed during prometaphase in control^RNAi^ cells. Bars, 5 μm. (H) Quantification of mitotic cells with null kinetochores from H. Error bars represent S.D. from three independent experiments. For each experiment, 50 mitotic cells were examined. (I) Immunofluorescence staining of MG132 + ZM treated mitotic HeLa cells depleted of endogenous Ndc80 and transfected with different Hec1-GFP constructs as indicated and stained for GFP (green), α-tubulin (red) with the chromosomes counterstained using DAPI. Bars, 5 μm. (J) Quantification of mitotic cells with null kinetochores from A. Error bars represent S.D. from three independent experiments. For each experiment, 200 mitotic cells were examined. ****P < 0.0001 (Student’s t test).

We then analyzed if the microtubule-binding mutants of Ndc80 could rescue the kinetochore-null phenotype produced in Ndc80^RNAi^ cells after ZM 447439 treatment. Our immunostaining data showed that cells rescued with Hec1^WT^ or nonphorylatable Hec1^9A^ mutant constructs rarely produced the kinetochore-null phenotype when Aurora B was inhibited (Fig. 4I, J, top two panels). In contrast, cells rescued with Hec1^9D^, Hec1^K166D^, or Hec1^LoopMut^ constructs showed a significantly high frequency of kinetochore-null phenotype in the same condition (Fig 4I, J, bottom three panels as indicated). This finding further supports the notion that the microtubule-binding activity of Ndc80 is directly responsible for the formation of initial kMT attachments during early mitosis.

### The Ndc80 Complex Coordinates with the Dynein Motor for Effective Kinetochore Capture

In addition to the Ndc80 complex, we tested if a similar kinetochore-null phenotype could be observed after the inhibition of dynein, CENP-E or Knl1, in combination with ZM447439 treatment. Interestingly, our immunofluorescence imaging data show that cells depleted of dynein in combination with ZM447439 treatment also showed a kinetochore null phenotype, but to a significantly lower extend as compared to Ndc80^RNAi^ cells. On the other hand, cells depleted of CENP-E or Knl1 did not exhibit a significant kinetochore-null phenotype under similar conditions (Fig 5A, B).

**Figure 5.**
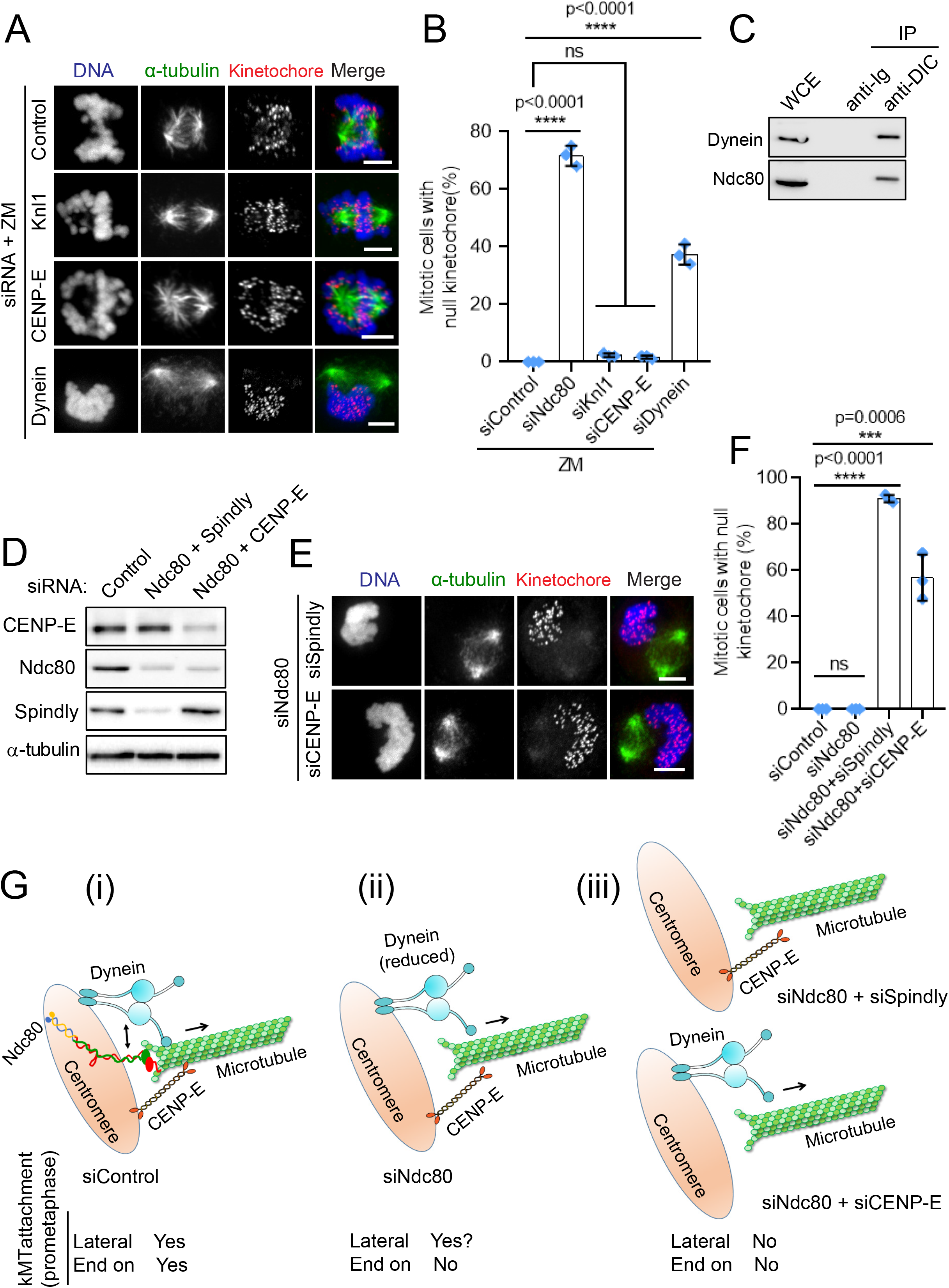
The Ndc80 complex coordinates with dynein for efficient kMT capture. (A) Immunofluorescence staining of MG132 + ZM treated mitotic HeLa cells depleted of the indicated proteins and stained for α-tubulin (green), a kinetochore marker (Zwint1 or CENP-A, red) with the chromosomes counterstained using DAPI. Bars, 5 μm. (B) Quantification of mitotic cells with null kinetochores from A. Error bars represent S.D. from three independent experiments. For each experiment, 200 mitotic cells were examined. ****P < 0.0001 (Student’s t test). (C) Control (Ig) and anti-dynein IC immunoprecipitations (IPs) were performed using whole cell extracts (WCE) prepared from nocodazole-treated prometaphase cells and the western blots of the IPs were probed with anti-dynein and anti-Ndc80 antibodies. (D) Western blot analysis of lysates obtained from HeLa cells co-depleted of Ndc80 + Spindly or Ndc80 + CENP-E as compared to control cell lysates. The blots were probed with antibodies against proteins that are indicated on the left. (E) Immunofluorescence staining of mitotic HeLa cells co-depleted of either Ndc80 + Spindly (top panel) or Ndc80 + CENP-E (bottom panel) and stained for α-tubulin (green), a kinetochore marker (Zwint1, red) with the chromosomes counterstained using DAPI. Bars, 5 μm. (F) Quantification of mitotic cells with null kinetochores from E. Error bars represent S.D. from three independent experiments. For each experiment, 200 mitotic cells were examined. ****P < 0.0001 (Student’s t test). (G) Cartoon depicting the coordinated action of Ndc80, Dynein and CENP-E in the formation of initial kMT attachments during early mitosis. During the kMT capture process, Ndc80 at the kinetochore is involved in attaching to the microtubules, not only via dynamic endon kMT contacts but also via dynamic lateral contacts. The kinetochore motors, dynein and CENP-E, assists Ndc80 by making lateral kMT contacts. In the absence of the Ndc80 complex, only about ~ 30% of lateral kMT contacts are made, mediated by dynein and CENP-E. Following the initial attachment stage, all kinetochores depend on dynein for spindle poleward motility, and event that is coordinated with the Ndc80 complex to constitute a processive kMT capture event. Loss of function of both the Ndc80 complex as well as the kinetochore motors is thus required for the complete inhibition of kMT capture.

The observation that the co-inhibition of Ndc80 and Aurora B resulted in a complete loss of initial kMT attachments was quite intriguing and we wanted to explore this phenotype further. It has previously been shown that Aurora B kinase activity is required to recruit the microtubule motors, dynein and CENP-E that are responsible for chromosome congression to the kinetochores (Ditchfield et al., 2003; Kasuboski et al., 2011; Murata-Hori et al., 2002). Moreover, our previous findings showed that Ndc80 competes with dynein to bind microtubules (Amin et al., 2018). We first tested if there is a physical association of Ndc80 with the two kinetochore microtubule motors in early mitosis by an immunoprecipitation assay. Our experiments showed that Ndc80 indeed interacts with dynein (Fig 5C) but not with CENP-E (data not shown) in prometaphase cell extracts. This result along with the observation that CENP-E inhibition did not produce significant defects in initial kMT attachments by itself or in combination with ZM447439 treatment, suggested to us that the kinetochore-null phenotype was likely produced due to improper recruitment of the dynein motor to Ndc80-depleted kinetochores after Aurora B inhibition. To test this further, we assessed dynein localization to prometaphase kinetochores in Ndc80-depleted cells treated with ZM447439. Even though we observed a minor reduction (~ 30%) in the immunostaining of dynein in non-nocodazole-treated, Ndc80-depleted kinetochores, substantial levels of kinetochore dynein were still retained in this case relative to control cells (Fig. S4D, top and middle panel; Fig S4E). However, we found that dynein immunostaining at kinetochores was almost completely lost (> 95%) when Ndc80-depletion was combined with Aurora B inhibition (Fig, S4D, bottom panel; Fig S4E).

To substantiate our results further, we performed co-depletion experiments of Ndc80 with either dynein or CENP-E. Since we observed that the co-depletion of the dynein motor and the Ndc80 complex was too toxic for the cells to tolerate, we depleted, Spindly, a recruiter of dynein to the kinetochore, in place of dynein. We confirmed the efficiency of depletion of the target proteins by immunoblotting (Fig 5D). Interestingly, we found the co-depletion of Ndc80 and Spindly, yielded an even higher frequency of the kinetochore-null phenotype (> 90%) compared to that observed in cells co-inhibited of Ndc80 and Aurora B (~75%) (Fig. 5E, F). Co-depletion of Ndc80 and CENP-E, on the other hand, also yielded a significant kinetochore-null phenotype than that observed with the depletion of either of the proteins individually, but weaker than that observed for Ndc80+Aurora B or Ndc80+Spindly co-inhibition (Fig. 5E, F and data not shown). Taken together, these data suggest that processive kMT capture during early mitosis depends on the concerted function of the both the Ndc80 complex and the dynein motor, and to a lesser extent, on CENP-E as well.

Overall, our study identifies a previously unreported function of the human Ndc80 complex during early mitosis. The Ndc80 complex was found to be localized to kinetochores during mitosis from prophase to anaphase (Ciferri et al., 2005; DeLuca et al., 2005; Varma et al., 2013). The main function of Ndc80 in stabilizing load-bearing kMT attachments in metaphase has been well studied. Its localization to kinetochores in early mitosis and strong affinity for binding to microtubules thus led us to test if the function of Ndc80 is required for kinetochore capture during early mitosis. It has been previously shown that chromosomes captured by spindle microtubules move poleward rapidly, via lateral kMT attachments and motility mediated by dynein (Rieder and Alexander, 1990; Vorozhko et al., 2008; Yang et al., 2007). *In vitro* analysis showed that microtubule-binding to kinetochores is reduced by dynein/dynactin inhibition (Vorozhko et al., 2008), and that kinetochore dynein prevents premature stabilization of load-bearling, end-on kMT attachments (Amin et al., 2018; Barisic and Maiato, 2015; Cheerambathur et al., 2013; Gassmann et al., 2008). Our data suggest that dynein, however, is not required for a large fraction of initial kMT attachments that are formed during early mitosis in Human cells (Fig S2C, D). This is likely to be especially true for those attachments involving the dynamic end-on contact of microtubule plus-ends with individual kinetochores during the ‘search and capture’ process in early mitosis that we surmise, are mediated by the Ndc80 complex. Further, our studies suggest that other kinetochore proteins including, dynein-dynactin, CLIP-170, Spindly and CENP-E, which localize to kinetochores and are important for chromosome alignment however do not possess a significant role in the initial kMT attachment process during early mitotic progression in normal cells (Fig S2C). These factors, however, are still likely to be required to form lateral kMT attachments with the microtubule lattice as has been shown by previous studies. Further, in the scenario where normal mitotic progression is severely delayed by defective kMT capture in Ndc80-inhibited cells, we predict that the above-mentioned microtubule-binding factors are also likely to compensate for the Ndc80 loss of function and rescue chromosome alignment to a substantial extend.

More importantly, in this study we show for the first time that the kinetochore-bound Ndc80 complex plays an essential role in initial kMT attachment formation during early mitosis in humans. As was just mentioned, many of these attachments are possibly dynamic end-on kMT contacts with mono-oriented kinetochores, which are thus critical for chromosome capture, leading to severe defects in kMT capture after the inhibition of the Ndc80 complex. While Ndc80 is likely to be unique in its ability to form dynamic end-on kMT attachments, the complex could also assist the kinetochore motors, Dynein and CENP-E, in forming lateral attachments during early mitosis. Original studies in yeast had found that the Ndc80 complex is important to from lateral kMT interactions during closed mitosis (Tanaka et al., 2005). As it is not clear if such a scenario exists during open mitosis in human cells and as our studies do not exclude this possibility, these points, taken together only amplify the key role played by Ndc80 in the kinetochore capture process. Here, we propose a model (Fig 5G) for efficient kMT capture during early mitosis mediated by the coordinated function of the Nc80 complex and dynein. In this model, kinetochore capture by microtubules during early mitosis could be further broken down into two successive steps. The initial step of kMT attachments, which we find is primarily mediated by the Ndc80 complex and the later step of poleward movement of the kinetochores during chromosome congression that has been established to primarily be driven by dynein.

As expected, the role of the Ndc80 complex in initial kMT attachments is dependent on its key microtubule-binding elements, the CH domain and the tail domain, at the N-terminus of the Hec1 subunit. The phosphorylation of these domains by Aurora B, possibly coupled with kinetochore phosphatase activity maintains a reduced affinity state of the Ndc80 complex for microtubules in prophase and prometaphase (Cheeseman et al., 2006; Foley and Kapoor, 2013). We propose that this version of the Ndc80 complex with the reduced affinity for microtubules is essential for initial kMT attachments during early mitosis. In support of this notion, the nonphosphorylatable Hec1^9A^ mutant showed efficient microtubule-binding to facilitate proper kMT capture similar to what is observed in cells rescued with Hec1^WT^. The phosphomimetic Hec1^9D^ mutant electrostatically impedes microtubules to bind with Ndc80 and thus does not allow to produce effective initial kMT attachments. Additionally, the Hec1^K166D^ mutant which has a single amino acid modification does not allow the formation of proper initial kMT attachments due to a conformational change in Hec1 that inhibits the CH domain binding to microtubules. Thus, as for load-bearing kMT attachments in metaphase, the initial kMT attachments are also regulated by Aurora B-mediated phosphoregulation of the N-terminal tail domain and the CH domain of the Hec1 subunit.

As mentioned earlier, it is worth noting that the Ndc80 complex could be instrumental in forming initial kMT attachments by two different mechanisms, both dependent on its ability to bind microtubules. The first mechanism, strongly supported by our observations is that the microtubule-binding activity of the complex is required to directly bind to *de novo* microtubules emerging from the spindle poles via the formation of lateral and/or end-on dynamic kMT contacts. The 2^nd^ equally plausible mechanism, the observation of which is limited by the resolution of our imaging technology is that the microtubule-binding activity of the Ndc80 complex could be essential for the formation of *de novo* microtubules originating at the kinetochores. Acentrosomal microtubules that are generated by such an Ndc80-mediated mechanism could it turn couple with the Augmin complex-dependent microtubule branching/bundling within the mitotic spindle (David et al., 2019; Luo et al., 2019), thus contributing to kinetochore capture and kMT maturation during early mitosis.

Recent studies have reported that the Ndc80-mediated stabilization of load-bearing kMT attachments prematurely during early mitosis is inhibited by kinetochore dynein (Amin et al., 2018; Cheerambathur et al., 2013; Gassmann et al., 2008). From our current findings we surmise that the function of Ndc80 complex in forming initial kMT attachments during early mitosis should precede the onset of inhibition of its function in kMT stabilization by dynein. While our experiments have failed to identify a role for other kinetochore factors apart from the Ndc80 complex in the formation of dynamic end-on kMT contacts during early mitosis, it is possible that, yet unknown mechanisms might also contribute to the capture process and hence to successful chromosome alignment. Once captured, dynein’s processive motor activity takes over to drive the kinetochores poleward for chromosome alignment. We propose a kinetochore ‘hand-over’ mechanism of coordination between these two microtubule-binding complexes, where a vast majority (~ 70 %) of the initial kMT attachments are facilitated by the Ndc80 complex which in turn hands-over the captured kinetochore to dynein for poleward kinetochore motility. This non-processive, but moderately robust kMT interactions in early mitosis facilitated by the Ndc80 complex would be critical to enhance the efficiency of kMT capture as prometaphase cells wait for the build-up of kinetochore dynein levels so that they can switch to dynamic kMT interactions and processive kinetochore motility. It is interesting to note that the Aurora B phosphorylated version of the Ndc80 complex, possibly still retains considerable affinity of microtubules to favor initial kMT contacts during early mitosis. Once the chromosomes attain proper alignment and biorientation, the Ndc80 complex at kinetochores serves its well-established function in the formation of load-bearing kMT attachments during metaphase and anaphase.

The additive nature of the Ndc80 and dynein co-inhibition phenotype in the context of initial kMT attachments suggest a possibly indirect mechanism of coordination between these two kinetochore complexes, in addition to their individual contributions to this process. It is however not clear at this point how the interaction we observe between dynein and the Ndc80 complex in prometaphase cells contribute to the coordinated function between these complexes for efficient kinetochore capture. The best explanation is that their interaction somehow contributes to the efficiency of the kinetochore ‘hand-over’ mechanism for processive capture. Another intriguing possibility is that the Ndc80 complex might be important to relieve the autoinhibition of the dynein motor that in turn could ‘prime’ the motor for processive minus-end directed motility. It has been proposed that the dynein motor is maintained in an auto-inhibited state prior to cargo-binding (Torisawa et al., 2014; Zhang et al., 2017). A 3^rd^ possibility is that the interaction contributes to the kinetochore-binding of at least a small fraction of dynein molecules. However, the severe defects in kinetochore capture observed after inhibiting Ndc80 as compared to the relatively minor phenotype obtained after dynein inhibition suggests that the possible loss of kinetochore dynein is likely not the main cause for defective capture after Ndc80 inhibition. Further studies are clearly required to delineate the molecular underpinnings of dynein-Ndc80 coordination during early mitosis.

## Materials and methods

### Cell culture, transfections, and drug treatments

HeLa and RPE1 cells were grown in Dulbecco’s modified Eagle medium (DMEM, Life Technologies) supplemented with 10% fetal bovine serum at 37 °C in humidified atmosphere with 5% CO_2_. The doxycycline (dox) inducible CRISPR/Cas9 Nuf2 knockout HeLa cell line (generous gift from Iain Cheeseman, MIT) was maintained in the absence of dox.

For RNA interference (RNAi) experiments, cells were transfected at 30-50% confluence using Dharmafect 2 (Dharmacon) according to the manufacturer’s instructions and analyzed 48-72 h after transfection. For rescue experiments, RNAi-refractory constructs were transfected into cells using Lipofectamine 3000 (Life Technologies) for 12 h followed by siRNA transfection. Cells were prepared for analysis after 48 h of siRNA transfection.

For testing kinetochore microtubule capture, cells were treated with STLC (5 μM) for 2 h prior to fixation to arrest them in early mitosis with monopolar mitotic spindles. Nocodazole (3 μM) was used to depolymerize the microtubule. To inhibit Aurora B kinase, 5 μM ZM 447439 (APExBio) was added to the medium, and cells were fixed after 1 h of incubation. Cells were treated with 10 μM MG132 to prevent mitotic exit after the inhibition of Aurora B kinase. Cas9 expression was induced by 1 μM doxycycline hyclate (Sigma).

siRNAs targeting 5’ UTR Ndc80 (DeLuca et al., 2011), 3’ UTR Dynein (Amin et al., 2018), and siRNAs targeting cDNAs for CLIP-170 and CENP-E (Amin et al., 2015), Nuf2 (DeLuca et al., 2002), Knl1 (Kiyomitsu et al., 2007) were used this study. The Smart Pool ON-TARGET plus siRNAs was used for Spindly knockdown (Amin et al., 2018). All siRNAs were used at 100 nM concentration except for dynein and CLIP-170 (200 nM).

### Antibodies

The primary monoclonal mouse antibodies used were anti-Hec1 9G3 (Abcam, ab3613) for IF at 1:400, and for WB at 1:1000; anti-Dynein Intermediate Chain (DIC) clone 74.1 (EMD Millipore, MAB1618) for IF at 1:300, for WB at 1:1000, anti-CENP-E 1H12 (Abcam, ab5093) IF at 1:400, and anti-α-tubulin DM1A (Santa Cruz, Sc32293) IF at 1:750, WB at 1:1000. The primary rabbit polyclonal antibodies used were anti-Hec1 (Bethyl Laboratories, A300-771A), anti-CLIP-170 (Santa Cruz, H-300) for IF at 1:750; anti-Zwint1 (Bethyl, A300-781A) for IF at 1:400; anti-Spindly (a gift from Dr. Reto Gassmann, UC San Diego) for IF at 1:5000, and for WB at 1:1000; anti-CENP-E (Boster, M04553-1) for WB at 1:1000. We also used a human anti-centromeric (Immunovision) at 1:500 for IF.

### Immunofluorescence

HeLa cells grown on a glass coverslip were fixed in cold methanol (−20°C) or 4% formaldehyde after pre-extraction with 0.1% Triton X-100 followed by blocking with 3% bovine serum albumin (BSA) in PBS and incubated with primary antibodies for 1 h at 37 °C followed by washing with PBS (137mM NaCl, 2.7mM KCl, 10mM Na_2_HPO_4_ and 1.8mM KH2PO_4_, pH 7.4) supplemented with 0.02% Triton X-100. The secondary antibodies coupled with Alexa-Fluor-488/647 or Rhodamine Red-X (Jackson ImmunoResearch Laboratories, Inc.) were used at a dilution of 1:250, and DNA was counterstained with 1 mg/ml DAPI.

For nocodazole washout assay, cells were treated with nocodazole (3 μM) for 3 h and transferred to DMEM medium at 37 °C after washing out nocodazole three times with PBS and once with warm DMEM medium. Cells were fixed at 0- and 10-min incubation with DMEM medium.

### Image acquisition and analysis

For image acquisition, three-dimensional stacks were obtained through the cell using a Nikon Eclipse TiE inverted microscope equipped with a Yokogawa CSU-X1 spinning disc, an Andor iXon Ultra888 EMCCD camera and an x60 or x100 1.4 NA Plan-Apochromatic DIC oil immersion objective (Nikon). For fixed cell experiments, images were acquired at room temperature as Z-stacks at 0.2 μm intervals controlled by NIS-elements software (Nikon). Images were processed in Fiji ImageJ and Adobe Photoshop CC 2018 and represent maximum-intensity projections of the required z-stacks.

For Live-cell imaging, HeLa cells stably expressing both mCherry H2B and GFP-α-tubulin or only GFP-H2B were cultured in 35mm glass-bottomed dishes (MatTek Corporation). Before 30 min of imaging, cell culture medium was changed to pre-warmed L-15 medium (Gibco) supplemented with 20% fetal bovine serum and 20mM HEPES, pH 7.0. Live-experiments were carried-out in an incubation chamber for microscopes (Tokai Hit Co., Ltd) at 37 °C and 5% CO_2_. Image recording was initiated immediately after adding MG132 (unless otherwise stated) using an x60 1.4 NA Plan-Apochromatic DIC oil immersion objective mounted on an inverted microscope (Nikon) equipped with an Andor iXon Ultra888 EMCCD camera or an Andor Zyla 4.2 plus sCMOS camera. Twelve 1.2 μm-separated z-planes covering the entire volume of the cell were collected at every 10 min up to 12 h.

To quantify the kinetochore motions, images were captured at 0.8 μm intervals every 5 s. Sister kinetochore pairs chosen for analysis were located at the periphery of the monopolar spindle. Kinetochore movements were tracked using GFP-CENP-A fluorescence using the manual-tracking tool in ImageJ software.

To measure the distance between kinetochore and nearest microtubule loci (attached or unattached), we manually selected a suitable image from a z stack carrying the concerned kinetochore and microtubule. We then calculated the distance of separation between the perceived center of the kinetochore and the end of the microtubule using ImageJ software.

All images and movies were processed in ImageJ/Fiji ImageJ and Adobe Photoshop CC 2017. All images represent maximum-intensity projections of the required z-stacks. Statistical tests performed are specified in figure legends.

### Western blotting

Cell lysates were prepared with lysis buffer [150 mM KCl, 75 mM HEPES of pH 7.5, 1.5 mM EGTA, 1.5 mM MgCl_2_, 10% glycerol, 0.1% NP-40, 30 mg/ml DNase, 30 mg/ml RNase, complete protease inhibitor cocktail (Roche) and complete phosphatase inhibitor cocktail (Sigma)]. Protein concentration of cell lysate was measured using the Coomassie protein assay kit (Thermo Scientific). Proteins were separated on SDS-PAGE, electroblotted onto a nitrocellulose blotting membrane (Amersham, GE Healthcare) and subjected to immunodetection using appropriate primary antibodies. Blocking and antibody incubations were performed in 5% non-fat dry milk. Proteins were visualized using horseradish peroxidase-conjugated secondary antibodies diluted at 1:2,000 (Amersham) and the ECL system, according to the manufacturer’s instructions (Thermo Scientific).

### Statistical analysis

The statistical analyses for scattered graphs were carried out using GraphPad Prism Software (version 8.1.0). Samples for analysis in each data set were acquired in the same experiment, and all samples were calculated at the same time for each data set. A two-sided t-test was used for comparison of average for bar graphs.

## Acknowledgements

The Authors would like to thank Drs. Iain Cheeseman for providing inducible CRISPR/Cas9 Nuf2 knockout HeLa cell line and Jennifer Deluca for the Hec1 mutant constructs. This work was supported by an NIGMS grant to DV (R01GM135391) and by start-up funds from Northwestern University.

## Abbreviations List

DIC: Dynein intermediate chain
kMT: kinetochore-microtubule
Ndc80: nuclear division cycle 80
CENP-E: Centrosome-associated protein E
ACA (CREST): anti-centromere antiserum
siRNA: small interfering RNA
RNAi: RNA interference
KMN: Knl1-Mis12-Ndc80
CH: calponin homology
STLC: S-Trityl-L-Cysteine
Hec1: Highly expressed in cancer protein 1
CLIP-170: CAP-GLY domain containing linker protein 170;

## Author’s Contributions

M.A.A. and D.V. designed and conceived the research. M.A performed all the experiments and data analyses and wrote the original draft. D.A.W. performed experiments for the revision. M.A, and D.V. contributed to editing the manuscript.

## Supplemental Figure legends

**Figure S1.**
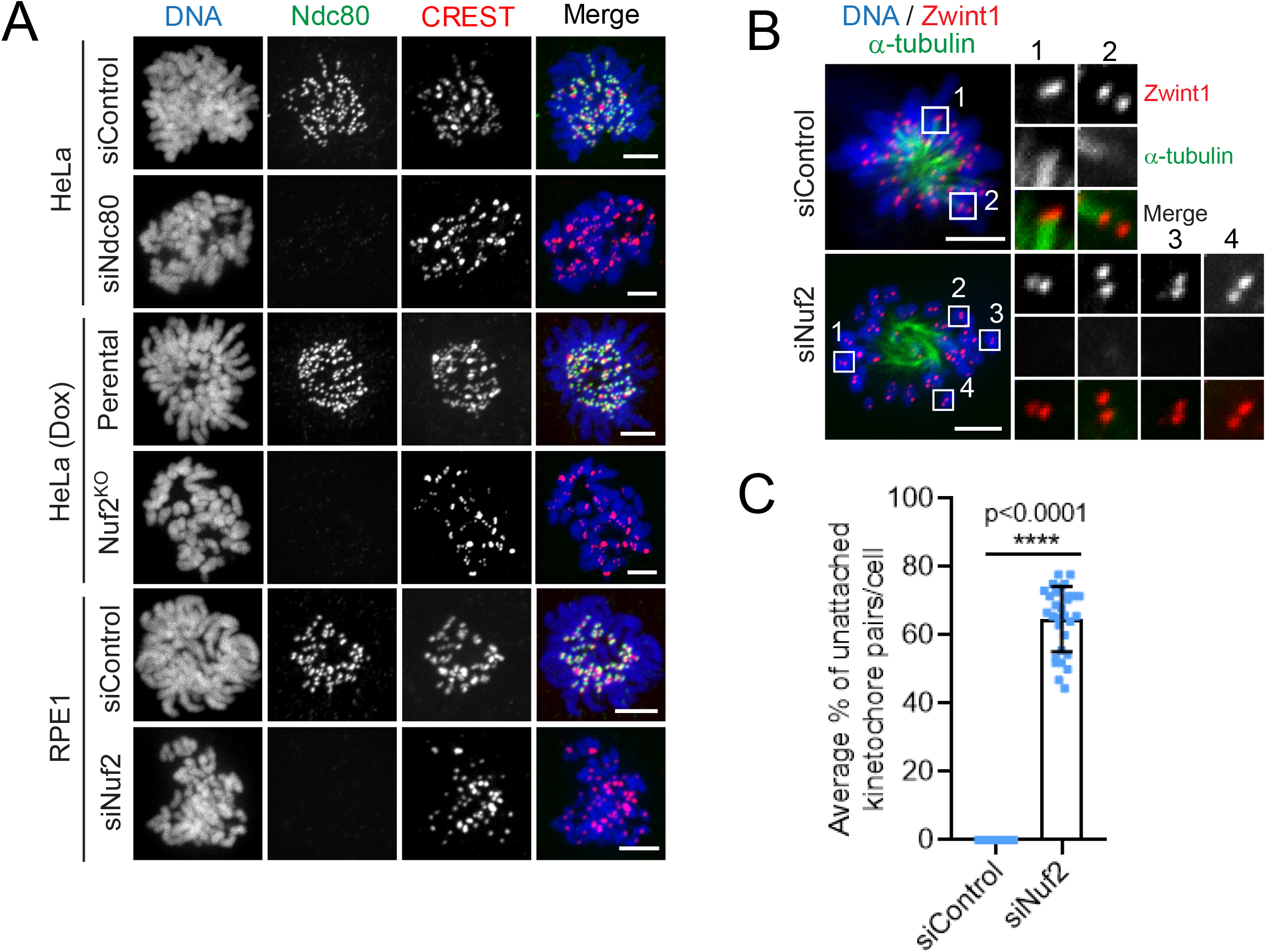
Immunofluorescence staining showing the loss of kinetochore Ndc80 and perturbed initial kMT attachments after different modes of Ndc80 inhibition. (A) Immunofluorescence staining of mitotic HeLa cells depleted of Ndc80 (top two panels) or knocked out of Nuf2 (middle two panels) or of RPE1 cells depleted of Nuf2 (bottom two panels) respectively in comparison to control or parental cells. The cells were stained for Ndc80 (green), a kinetochore marker CREST (red) with the chromosomes counterstained using DAPI. Bars, 5 μm. (B) Immunofluorescence staining of STLC-treated mitotic HeLa cells knocked out of Nuf2 (bottom panel) in comparison to parental cells (top panel) and stained for α-tubulin (green), a kinetochore marker Zwint1 (red) with the chromosomes counterstained using DAPI. Bars, 5 μm. Inset shows the kMT attachment status of individual kinetochore pairsafter Nuf2 KO. (C) Quantification of the status of kMT attachments in cells from B. Error bars represent S.D. from three independent experiments. For each experiment, 10 monopolar cells were examined. ****P < 0.0001 (Student’s t test).

**Figure S2.**
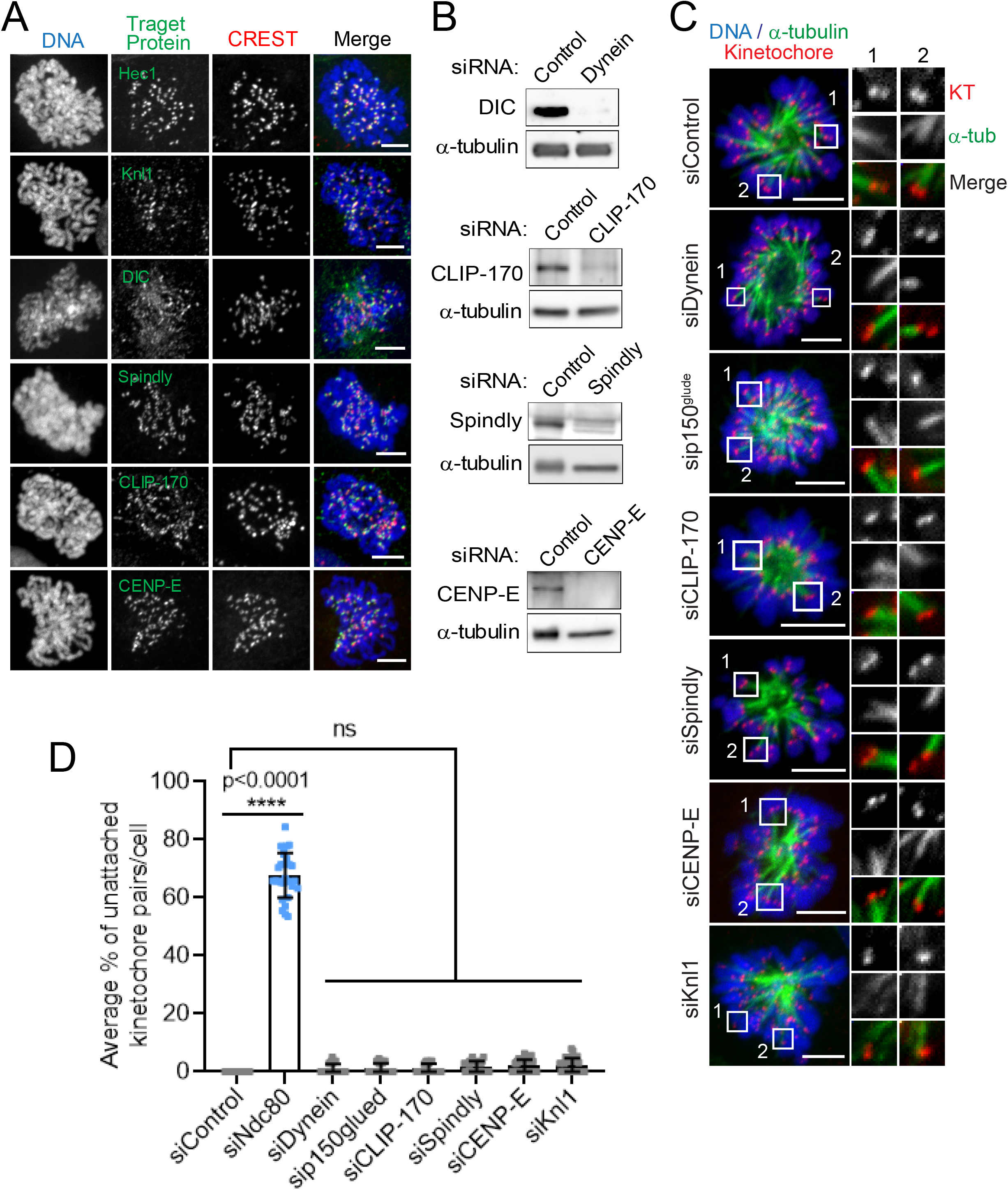
Localization of different kinetochore proteins in early prometaphase and analysis of their role in initial kMT capture. (A) Immunofluorescence staining of early prometaphase HeLa cells with different kinetochore proteins as indicated (green), and a kinetochore marker CREST (red) with the chromosomes counterstained using DAPI. Bars, 5 μm. (B) Western blot analysis of HeLa cells treated with indicated siRNAs. α-tubulin was used as a loading control. (C) Immunofluorescence staining of STLC-treated mitotic prometaphase cells depleted of the indicated target proteins as compared to control HeLa cells, and stained for α-tubulin (green), a kinetochore marker Zwint1 or CENP-A (red) with the chromosomes counterstained using DAPI. Bars, 5 μm. Inset shows the kMT attachment status of individual kinetochore pairs in the conditions indicated. (D) Quantification of the status of kMT attachments in cells from C. Error bars represent S.D. from three independent experiments. For each experiment, on an average ~ 20 kinetochore pairs from 10 monopolar cells were examined. ****P < 0.0001 (Student’s t test).

**Figure S3.**
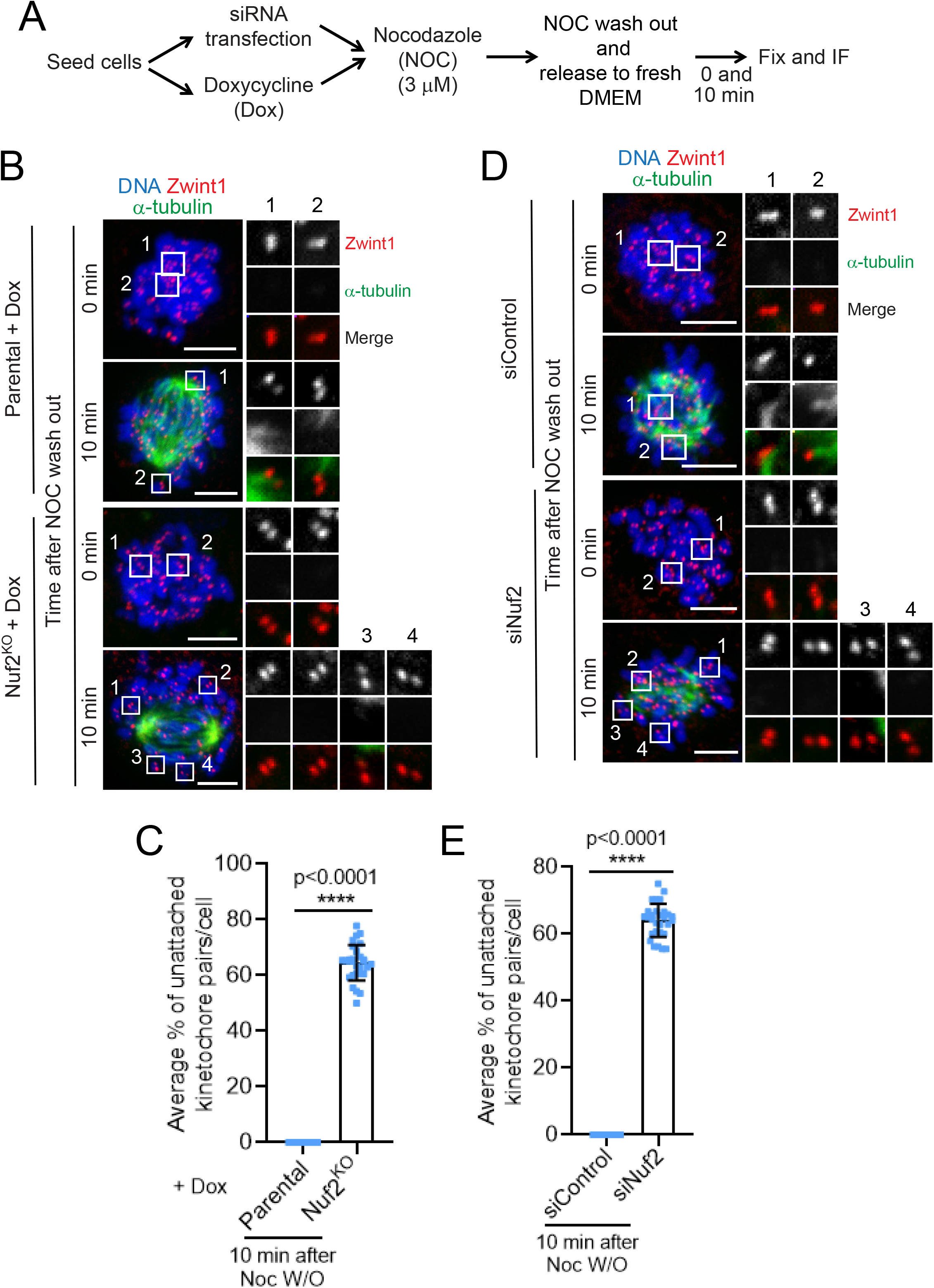
Nocodazole wash out assay to assess kinetochore capture by de novo microtubules early in mitosis after different modes of Ndc80 inhibition. (A) Cells were subjected to the indicated perturbation, treated with the indicated drug, and fixed according to the scheme. (B and D) Immunofluorescence staining of mitotic HeLa cells knocked out of Nuf2 (B) or of mitotic RPE1 cells depleted of Nuf2 (D) in comparison to control or parental cells respectively and immunostained for α-tubulin (green), a kinetochore marker Zwint1 (red) with the chromosomes counterstained using DAPI. Bars, 5 μm. Inset shows the kMT attachment status of individual kinetochore pairs in the indicated conditions. (C and E) Quantification of the status of kMT attachments in cells from B and D. Error bars represent S.D. from three independent experiments. For each experiment, 10 mitotic cells were examined. ****P < 0.0001 (Student’s t test).

**Figure S4.**
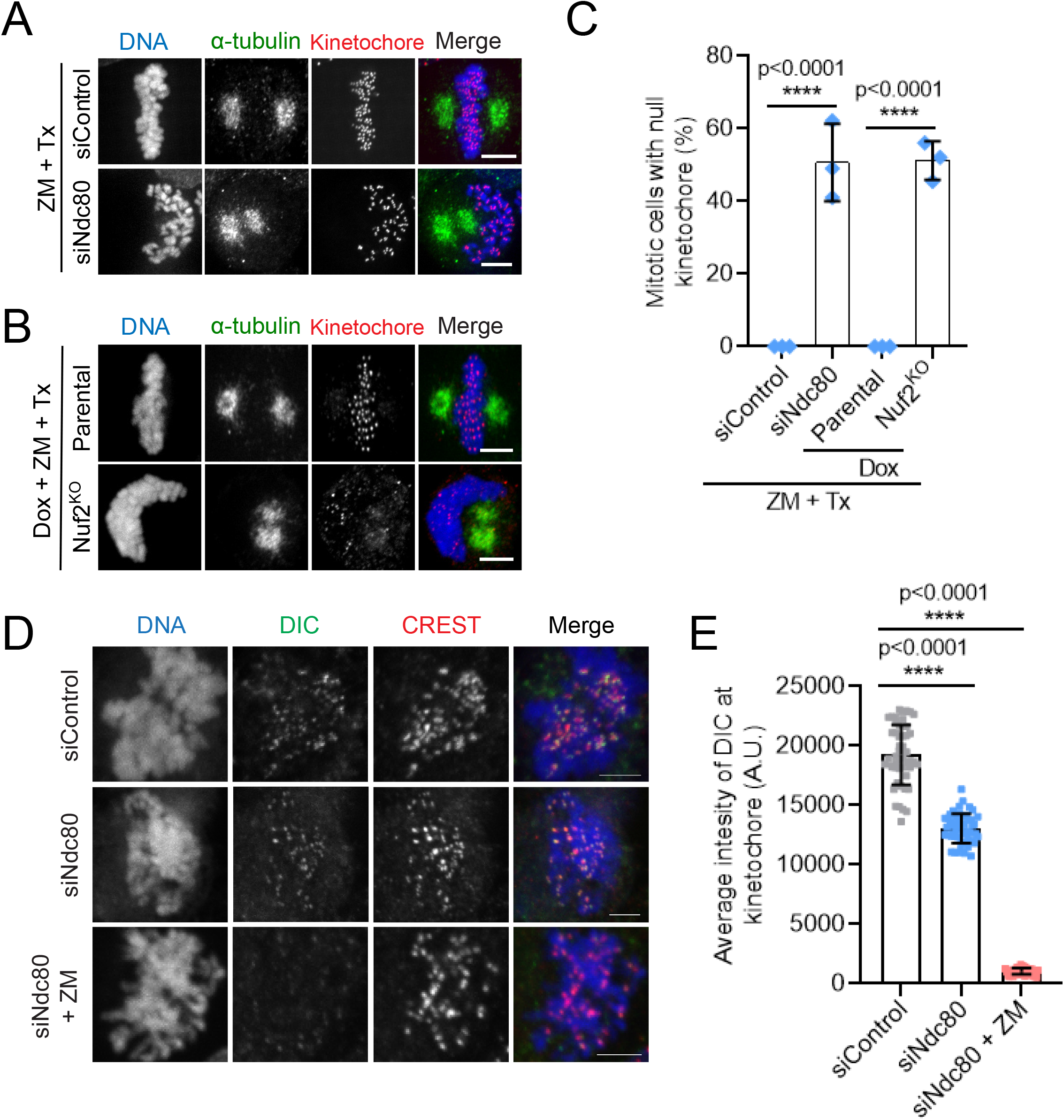
Further characterization of the kinetochore null phenotype and dynein kinetochore localization in Ndc80-inhibited cells. (A and B) Mitotic HeLa cells depleted of Ndc80 (A, bottom panel) or knocked out of Nuf2 (B, bottom panel) as compared to control^RNAi^ (A, top panel) or parental control cells (B, bottom panel), and followed by the indicated drug treatments, were immunostained for α-tubulin (green), a kinetochore marker Zwint1 or CENPA (red) with the chromosomes counterstained using DAPI. Bars, 5 μm. (C) Quantification of the frequency of mitotic cells with null kinetochores (see main text for more details) in samples from A and B. Error bars represent S.D. from three independent experiments. For each experiment, 200 mitotic cells were examined. ****P < 0.0001 (Student’s t test). (D) Analyses of dynein localization to kinetochores in Ndc80-depleted cells with or without Aurora B inhibition. Prometaphase HeLa cells were either treated with Ndc80 siRNA alone (middle panel) or in combination with ZM (bottom panel) as compared to untreated controls (top panel), followed by fixation and immunostaining for the dynein intermediate chain (DIC, green), a kinetochore marker (CREST, red) with the chromosomes counterstained using DAPI. Bars, 5 μm. (E) Quantification of data in D. A total of 50 kinetochores were analyzed from at least 5 different prometaphase cells.

## Supplemental Movie legends

**Movie S1.** Video of the dynamics of kinetochore pairs in HeLa cells stable expressing GFP-CENPA and GFP-Tubulin and treated with control siRNA. 0.6 um z sections were acquired at 15 sec intervals. Bars, 5 μm. Movie speed is 5 frames per second.

**Movie S2.** Video of the dynamics of kinetochore pairs in HeLa cells stable expressing GFP-CENPA and GFP-Tubulin and treated with siRNA for Ndc80. 0.6 um z sections were acquired at 15 sec intervals. Bars, 5 μm. Movie speed is 5 frames per second.

**Movie S3.** Video of kinetochore capture by microtubules during early prometaphase stage in HeLa cells stably expressing H2B-mCherry, GFP-α-tubulin, GFP-CENP-A and treated with control siRNA. Images were captured starting from the nuclear envelope breakdown until the time when chromosome alignment was attained. 0.6 um z sections were acquired at 1 min intervals. Bars, 5 μm. Movie speed is 5 frames per second.

**Movie S4.** Video of kinetochore capture by microtubules during early prometaphase stage of HeLa cells stably expressing H2B-mCherry, GFP-α-tubulin, GFP-CENP-A and treated with siRNA for Ndc80. Images were captured starting from the nuclear envelope breakdown until the time when complete chromosome alignment was expected to be attained (for around 30 min). 0.6 um z sections were acquired at 1 min intervals. Bars, 5 μm. Movie speed is 5 frames per second.

**Movie S5.** Video of chromosome congression during early prometaphase (where initial kMT attachment occurs) in parental HeLa cells treated with Hoechst to visualize the chromosomes. Images were captured starting from the nuclear envelope breakdown until the time when complete chromosome alignment was attained. 1 um z sections were acquired at 2 min intervals. Bars, 5 μm. Movie speed is 5 frames per second.

**Movie S6.** Video of chromosome congression during early prometaphase where initial kMT attachment occurs of CRISPR/Cas9 Nuf2^KO^ HeLa cells treated with Hoechst to visualize the chromosomes. Images were captured starting from the nuclear envelope breakdown until the time when complete chromosome alignment was expected to be attained (for around 30 min). 1 um z sections were acquired at 2 min intervals. Bars, 5 μm. Movie speed is 5 frames per second.

**Movie S7.** Video showing the behavior of kinetochores in HeLa cells expressing H2B-mCherry, GFP-α-tubulin and treated with control siRNA. Imaging was started after 30 min of adding MG132 and ZM. For the particular mitotic cell shown, images were captured starting from the nuclear envelope breakdown until the time when chromosomes were expected to align at the metaphase plate (for around 30 min). 0.6 um z sections were acquired at 2 min intervals. Bars, 5 μm. Movie speed is 5 frames per second.

**Movie S8.** Video demonstrating the kinetochore-null phenotype of HeLa cells expressing H2B-mCherry, GFP-α-tubulin and treated with siRNA for Ndc80. Imaging was started after 30 min of adding MG132 and ZM. For the particular mitotic cell shown, images were captured starting from the nuclear envelope breakdown until the time when chromosomes were expected to align at the metaphase plate (for around 30 min). 0.6 um z sections were acquired at 2 min intervals. Bars, 5 μm. Movie speed is 5 frames per second.

